# Exploring the potential of *Celtis aetnensis compounds* through Molecular Docking for Hepatocellular carcinoma

**DOI:** 10.1101/2024.02.28.582503

**Authors:** K.N. Kavitha, K. Revathi, Tamilamban Tamiraikani

**Author notes:** Corresponding author Dr. Kavitha. K.N., Meenakshi Academy of Higher Education and research, K K Nagar, Chennai-78., Ph;91-9384675125.

## Abstract

Hepatocellular carcinoma (HCC) accounts for 1% of the liver malignances; HCC progresses in hepatocytes primarily as a result of inflammation, oxidative stress, and primary liver disease. Thanks to the in-silico methods that help in the identification of the targets and the drugs/potential drugs that can inhibit these proteins that cater to the growth of the cancer cells. Plant actives have been long considered as potential sources of anticancer drugs. In this regard, we studied the actives from *Celtis tournefortii Lam (Celtis aetnensis) using* GCMS analysis. The protein structures of three receptors—NF-B P50 homodimer, FGF receptor 4, and vascular endothelial growth factor receptor 2 (VEGFR2)—were then subjected to docking studies using phytocompounds from *Celtis aetnensis*. Three receptor proteins were used in the docking analysis against 7 ligands. In this study, the PDB was used to obtain the structures of 3 cancer-related receptor proteins. The proteins were produced by eliminating water molecules, ligands using PyMol, and were then exposed to docking experiments. Docking studies revealed that the compounds with a binding energy ranging from -1.83 kcal/mol to -6.12 kcal/mol. The docking results revealed that eicosatetraenoic acid has a binding energy of-6.12 kcal/mol against the FGFR4 receptor. Sulfadoxine’s binding energy to the VEGFR2 receptor protein was-4.9 kcal/mol. Promethazine sulfoxide docks with an energy of -5.36 kcal/mol against NFKB P50. By attaching to the protein, these substances demonstrated good inhibitory activity. The results are supportive that these compounds may be used to treat HCC.

## INTRODUCTION

Hepatocellular Carcinoma (HCC) is a common form of liver malignancy responsible for approximately one million deaths worldwide annually (Mittal and El-Serag, 2013). The primary risk factors associated with HCC include chronic infection with hepatitis B virus (HBV) and hepatitis C virus (HCV), as well as exposure to aflatoxins, alcohol abuse and non-alcoholic fatty liver (Caillot et al., 2009). Chemotherapy drugs used in the treatment of HCC induces side effects in the nervous system. Therefore, a medication without side effects to be developed as a novel drug for the treatment of hepatocellular carcinoma.

*Celtis tournefortii Lam (Celtis aetnensis)* belongs to the genus Celtis with 70 species of shrubs widely present in the Northern Hemisphere. This woody plant has heart-shaped leaves with crenate on the edge. The plant has many traditional uses for the treatment of Gastrointestinal infections and to treat oral cavity infections which attribute to the medicinal values of the bioactive compounds like flavonoids, tannins and saponins. Numerous antioxidant compounds have been under scrutiny for their potential as chemo-preventive agents. The current study aims to explore the *in vivo* protective effects of natural compounds found in *Celtis tournefortii Lam (Celtis aetnensis)* against oxidative damage, particularly in relation to human cancer (Acquaviva, Rosaria, et al.,2016).

The growing interest in natural medicinal sources has led to the investigation of traditional compounds as potential treatments for various diseases (Sudha et al.2008). Virtual screening, utilizing molecular databanks, has proven to be effective method for the initial search of compounds with promising properties (Vyas et al.,2008). In the modern drug discovery process, molecular docking has emerged as a crucial research tool (Kitchen et al.,2004)

This study deals with molecular interactions of potential compounds of *Celtis tournefortii Lam (Celtis aetnensis)* against the virulence factors of hepatocellular carcinoma.

### MATERIALS AND METHODS

Seven anti-cancer compounds extracted from *Celtis aetnensis* were subjected to docking tests against the nuclear factor kappa B (NF-B) P50 homodimer, fibroblast growth factor (FGF) receptor 4, and vascular endothelial growth factor receptor 2 (VEGFR2). The three-dimensional structures of the receptor proteins were retrieved from Protein Data Bank (PDB). The PDFB IDs are 1SVC (NFKB P50), 4XCU (FGFR4), and 4ASD (VEGFR2). The ligand’s structures were acquired from PubChem. The active site of the proteins was retrieved from PDBsum. The compounds reported in *Celtis aetnensis* is given in Table 1

**Table 1:**
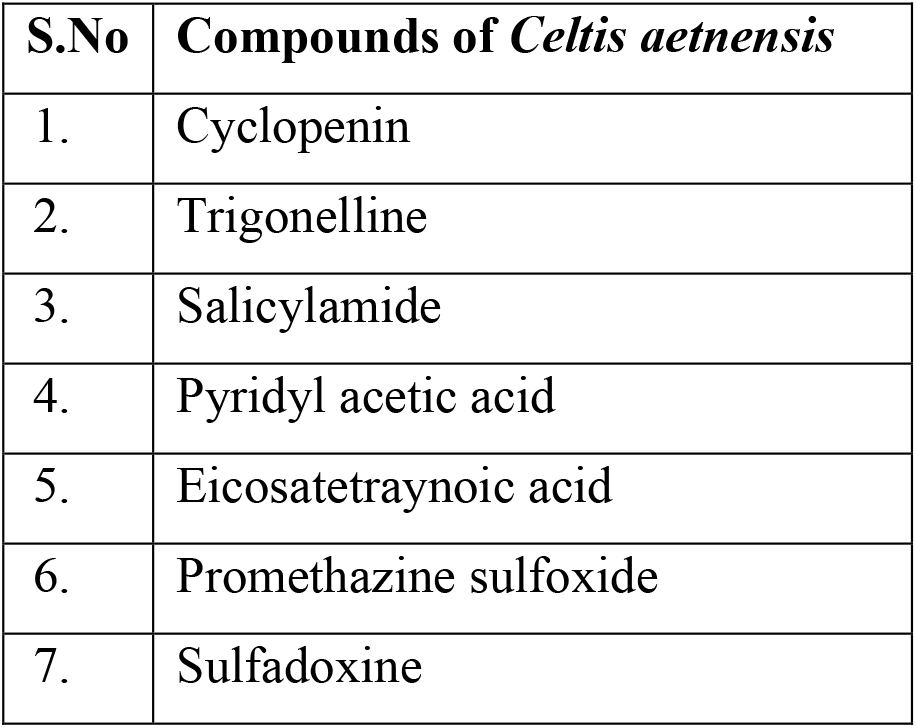
Compounds present in *Celtis aetnensis* used for docking study.

| S.No | Compounds of *Celtis aetnensis* |
| --- | --- |
| 1. | Cyclopenin |
| 2. | Trigonelline |
| 3. | Salicylamide |
| 4. | Pyridyl acetic acid |
| 5. | Eicosatetraynoic acid |
| 6. | Promethazine sulfoxide |
| 7. | Sulfadoxine |

### Molecular Docking analysis

Docking is a computational scoring method used to identify the optimal fit between a receptor and a ligand. Prior to docking, macromolecules were prepared by adding polar hydrogen and Kollman charges. A grid box was created to encompass the entire protein, and within this grid, calculations were performed for the atoms present. During docking, the ligand explores and identifies a binding cavity, and the of resulting macromolecule-ligand complex is computed. Numbers of such bound confirmations with stable energies were given as output. Those confirmations were analyzed and found in which of the bound confirmation the ligand is bound to the active site region. The conformational energy was noted. The binding efficiency was determined by the lower energy values obtained (Samant and Javle, 2020).

Molecular docking studies were performed by Auto Dock 4.2. which is based on Lamarckian Genetic Algorithm. Auto Dock uses two methods to perform docking such as energy evaluation based on grid and torsional freedom search. For docking experiment, the initial protein coordinates were extracted from the pdb file, and the ligand pdb files obtained from the PRODRG server were then utilized for docking studies using Auto Dock 4.2. Subsequently, the binding energy was calculated for each ligand, and contact analysis of the docked complexes was performed using Discovery Studio Visualizer 3.1.

### Discovery Studio Visualizer 3.1

Discovery Studio Visualizer is a freely available software utilized for conducting simulating and molecular modeling experiments for small and macromolecule. It is the product of Accelrys and has a wide use in academic and commercial industries particularly in biotechnology and pharmaceutical industries.Chu.et.al (2018) reported that Discovery Studio Visualizer generates 3D and 2D receptor-ligand interaction plots and enables the analysis of ligand binding pattens between the receptor protein and the ligand of interest.

## RESULT AND DISCUSSION

### Molecular docking studies

Seven potent anti-cancer compounds extracted from *Celtis aetnensis* were then put through docking tests against the nuclear factor kappa B (NF-B) P50 homodimer, fibroblast growth factor (FGF) receptor 4, and vascular endothelial growth factor receptor 2 (VEGFR2). Numerous cancers involve the FGFR4, VEGFR2, and NFKB P50 signaling pathways, and inhibiting their activity can greatly slow the growth of cancer. According to reports, the tyrosine kinase receptor (FGFR4) stimulates tumor cell growth, differentiation, and angiogenesis as well as increases the tumor cells’ capacity to withstand drugs (Liu et al., 2021). Due to their involvement in angiogenesis, which improves the delivery of nutrients and oxygen to the cancer cell, VEGFR2 is a key target in a variety of cancers (Lian et al., 2019). This enhances cancer cell proliferation, survival, and metastasis. By activating anti-apoptotic genes such TNFA, IL6, BCL2, and BCLXL, the NF-B P50 signaling pathway promotes tumor cell proliferation, angiogenesis, metastasis, and enhances cancer cell survival (Xia et al., 2018). Additionally, the NF-B P50 improves the immunosuppressive environment by enhancing macrophage M1 suppression (Xia et al., 2014).

Protein Data Bank (https://www.rcsb.org/) provided the three-dimensional structures of the receptor proteins. The PDFB IDs are 1SVC (NFKB P50), 4XCU (FGFR4), and 4ASD (VEGFR2). The ligand’s structures were acquired from PubChem. Using PDBSum, the active site area of FGFR4, VEGFR2 was discovered (Figure 1). The NFKB P50 structure 1SVC’s active site was not available in PDBSum. Hence the information about the active site amino acids Lys52, Ser243, Asp274, Lys, 275 was obtained from previous studies (Mukund et al., 2019)

**Figure 1.**
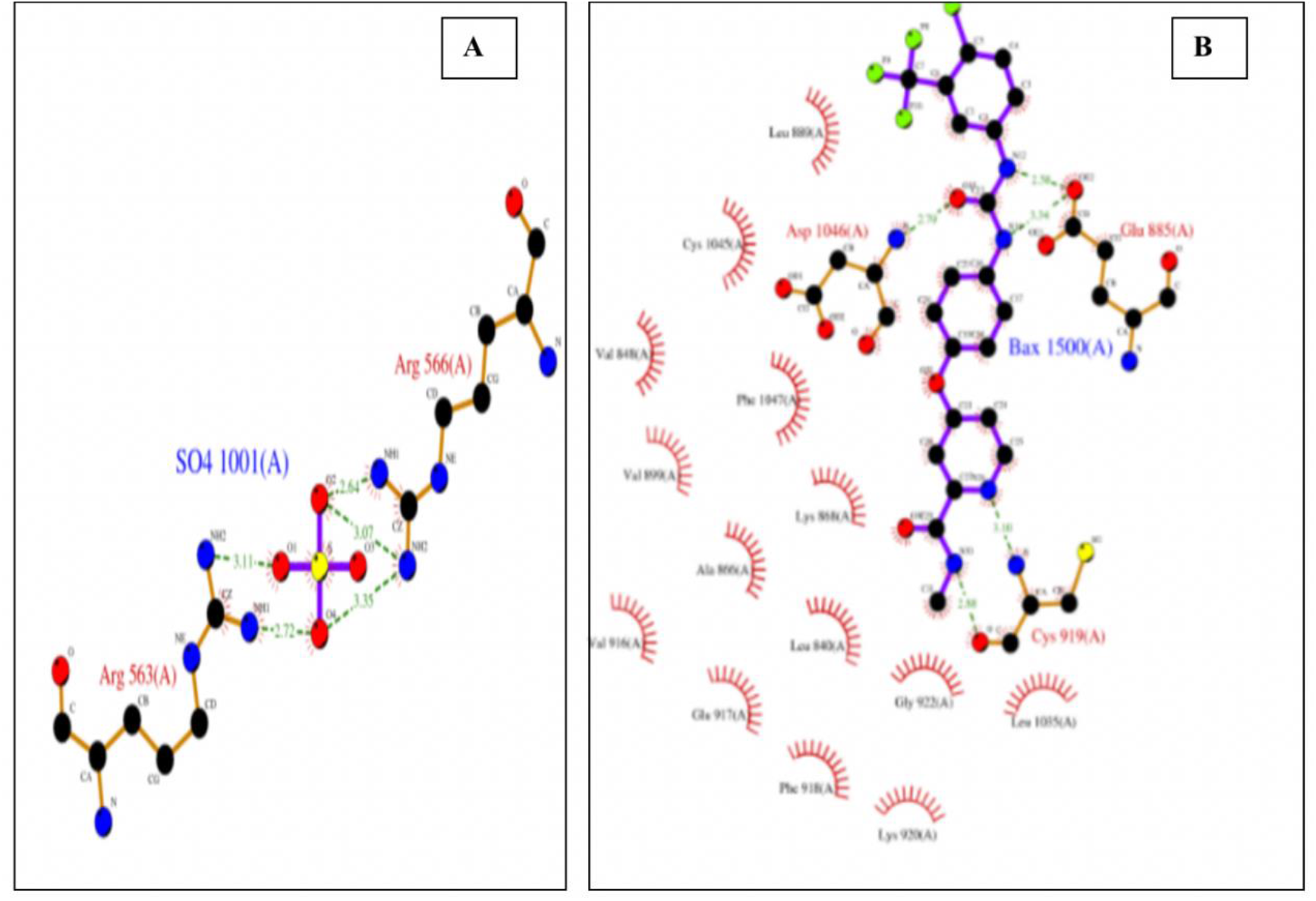
: Active site amino acids of A) 4XCU and B) 4ASD obtained from PDBsu.

The goal of molecular docking research is to identify prospective drugs that have a high affinity for the FGFR4, VEGFR2, and NFKB P50. The compound name, their hydrogen bonds, hydrophobic interactions and dock score (kcal/mol) are tabulated in Table.

Eicosatetraenoic acid made four hydrogen bonds with HIS638, ASP641, and LYS644 while binding to 4XCU with the least binding energy (−6.12 kcal/mol) of the other molecules. Trigonelline, which had a docking score of -5.94 kcal/mol and established hydrogen bonds with LYS644, ASP641, and HIS638 after that, was in second place. Sulfadoxine had a binding score of -5.89 kcal/mol with 4XCU, Promethazine Sulfoxide showed a binding score of -5.24 kcal/mol, Salicylamide had a binding score of -4.93 kcal/mol, Cyclopenin had a binding score of -4.9 kcal/mol, and Pyridyl Acetic Acid had a binding score of -4.71 kcal/mol (Figure 2).

**Figure 2.**
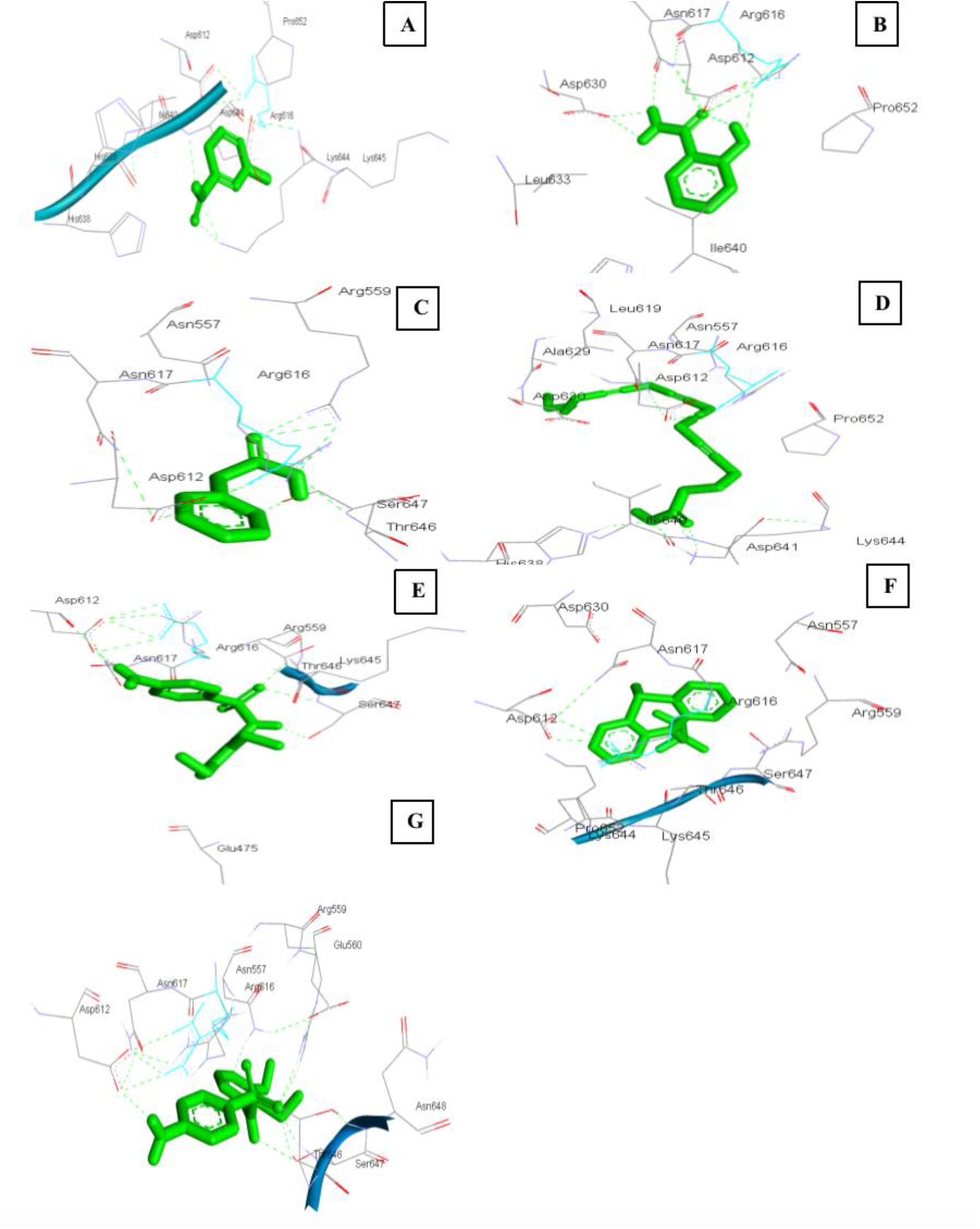
: Binding interactions of the protein fibroblast growth factor (FGF) receptor 4 (PDB id: 4XCU) with (a) Trigonelline, (b) Salicylamide, (c) Pyridyl acetic acid, (d) Eicosatetraynoic acid, (e) Cyclopenin, (f) Promethazine sulfoxide and (g) Sulfadoxine.

For the protein 1SVC, Promethazine sulfoxide displayed a least binding energy of -5.36 kcal/mol. Promethazine sulfoxide, however, did not create any hydrogen bonds with the receptor protein. Cyclopenin established a hydrogen bond with the amino acids ILE283, LYS336, and displayed a dock score of -5.32 kcal/mol. Docking scores of 4.87 kcal/mol, 4.76 kcal/mol, -4.3 kcal/mol, -4.18 kcal/mol, and -3.62 kcal/mol were found for the molecules eicosatetraenoic acid, pyridyl acetic acid, sulfadoxine, trigonelline, and salicylamide, respectively (Figure 3).

**Figure 3.**
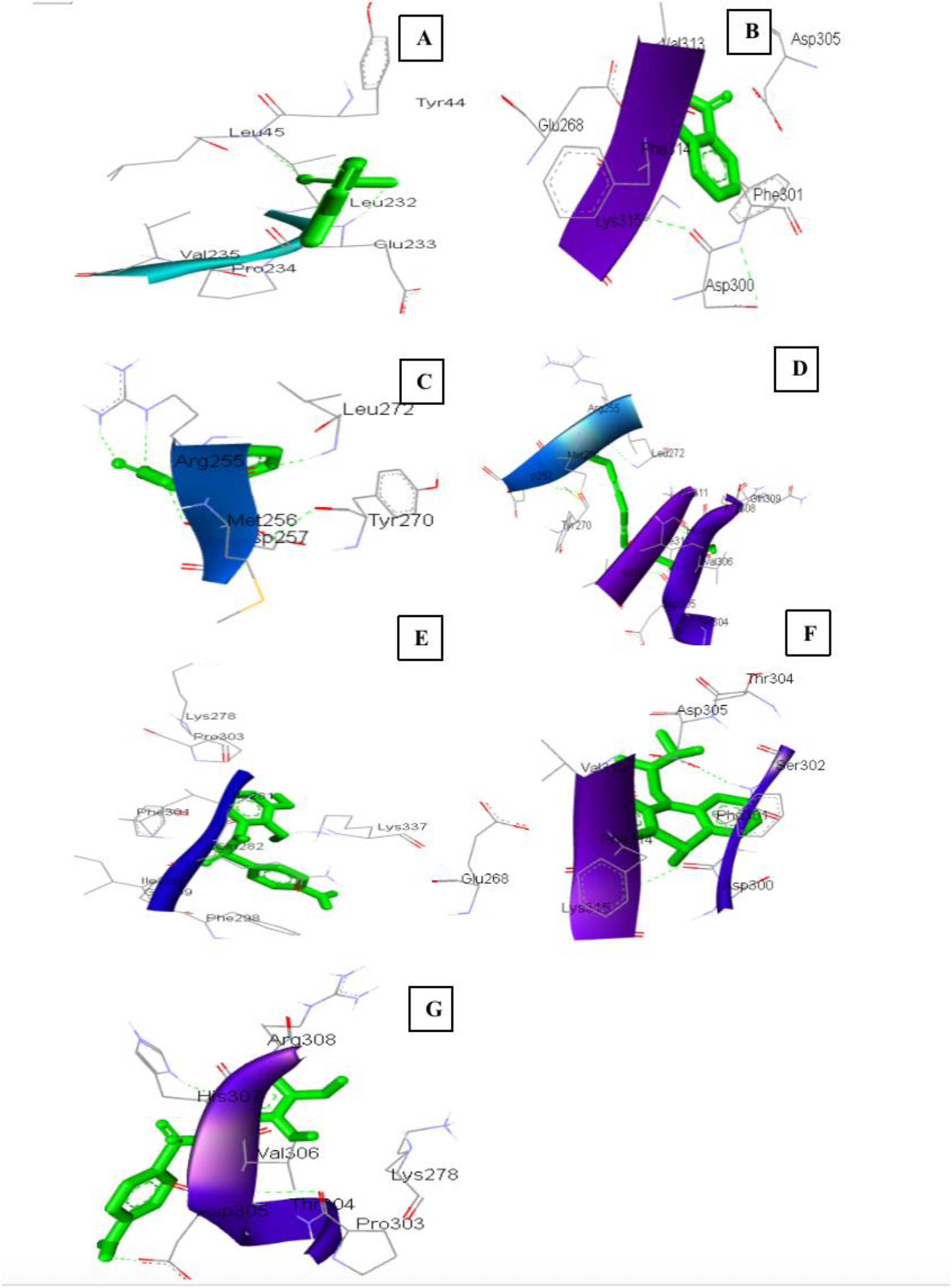
: Binding interactions of the protein fibroblast growth factor (FGF) receptor 4 (PDB id: 1SVC) with (a) Trigonelline, (b) Salicylamide, (c) Pyridyl acetic acid, (d) Eicosatetraynoic acid, (e) Cyclopenin, (f) Promethazine sulfoxide and (g) Sulfadoxine.

For 4ASD, Sulfadoxine made three hydrogen bonds with ARG932, SER1104, and ALA112 and had a binding score of -4.9 kcal/mol. The amino acids ARG932, SER1104, and ALA1127 made a hydrogen bond with cyclopenin and had a dock score of -4.86 kcal/mol. The substances promethazine sulfoxide, trigonelline, salicylamide, and pyridyl acetic acid showed dock scores of -4.8 kcal/mol, -3.89 kcal/mol, -4.41 kcal/mol, and -3.88 kcal/mol, respectively, with 4ASD (Figure 4).

**Figure 4.**
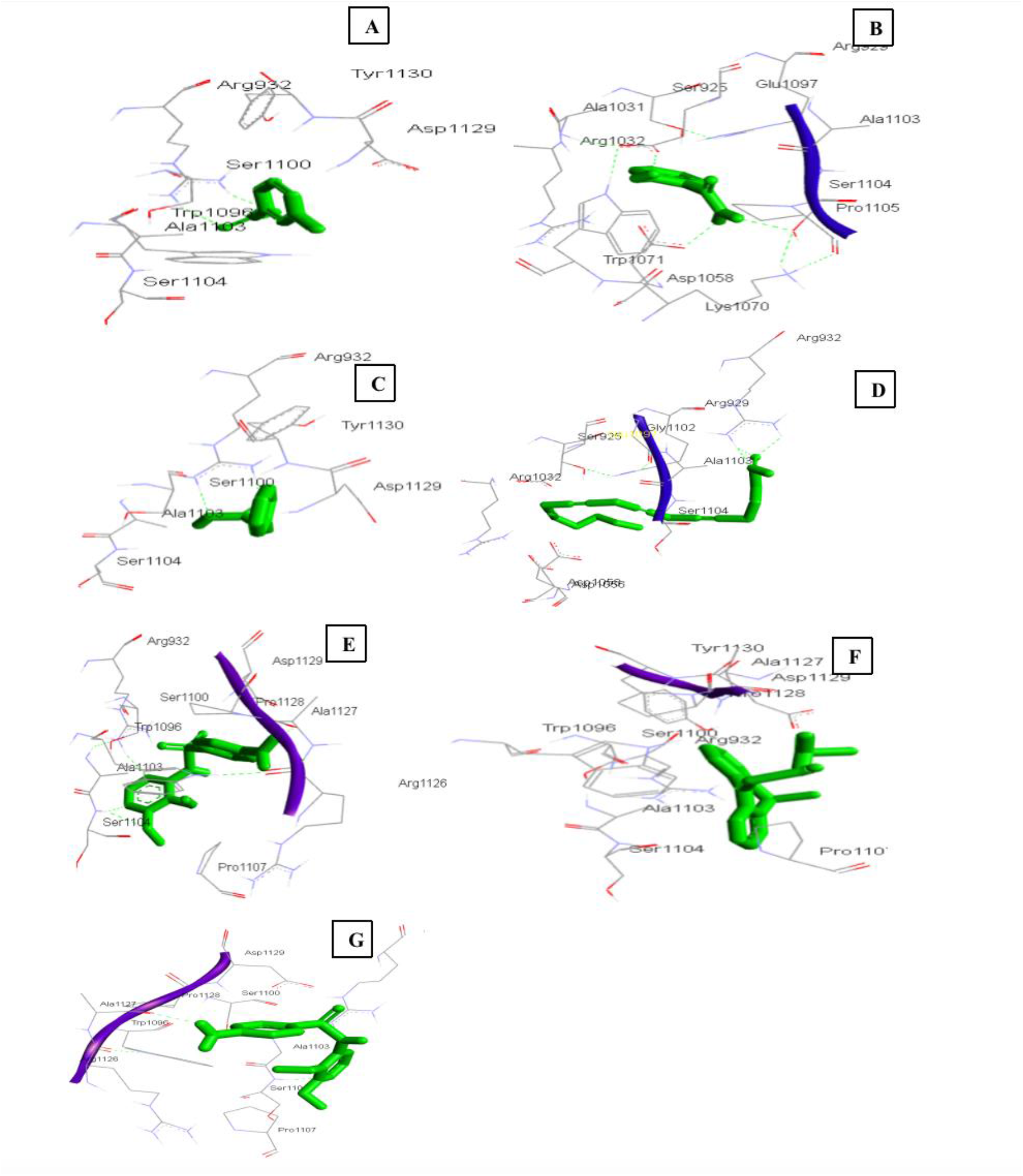
: Binding interactions of the protein fibroblast growth factor (FGF) receptor 4 (PDB id: 4ASD) with (a) Trigonelline, (b) Salicylamide, (c) Pyridyl acetic acid, (d) Eicosatetraynoic acid, (e) Cyclopenin, (f) Promethazine sulfoxide and (g) Sulfadoxine.

Binding affinity score of ligands with protein serves a crucial role in finding out the mechanism of pharmacological action of ligands. Inhibition of FGFR4, VEGFR2 and NFKB P50 signaling pathways can downregulate the proteins involved in cancer progression and metastasis. Compounds from *Cocculus hirsutus* showed good inhibition against the virulence factors of hepatocellular carcinoma (Thavamani et al., 2016). About 41 Phyto triterpenes originated from Vietnamese plants subjected to docking study proved to be promising against Mortalin inhibition activity. In a recent study by Song et al., (2022), Interleukin-1 receptor-associated kinase 1 was subjected to docking studies against natural compounds for hepatocellular carcinoma. In a recent thorough investigation by Mustafa et al. (2023), 1000 different plant phytochemicals were docked to HCC-related proteins. In order to investigate the compounds’ potential for inhibition, they were docked to the active site amino acids of the receptor proteins caspase-9 and the epidermal growth factor receptor. Based on their binding affinities and root-mean square deviation values, the top five compounds against each receptor protein were investigated as possible therapeutic candidates. As a result of this study, liquoric acid and limonin, were identified as potential drugs for the treatment of HCC in the future.

### CONCLUSION

The molecular docking of compounds from *Celtis aetnensis* clearly shows the manner of binding, participating amino acids, and the number of hydrogen bonds and other binding interactions. Hence these compounds could be subjected to experimental work against hepatocellular carcinoma.

**Table 1:** Docking Score, Hydrogen bond interactions of Protein and Ligand.

| **Protein** | **Ligand** | **Score**<br>**(kcal/mol)** | **No. of**<br>**hydrogen**<br>**bond** | **Hydrogen bond** |
| --- | --- | --- | --- | --- |
| 4XCU | Cyclopenin | -4.9 | 2 | ARG559, SER647 |
| 4XCU | Trigonelline | -5.94 | 4 | LYS644, ASP641, HOS638 |
| 4XCU | Salicylamide | -4.93 | 3 | ASP612, ASN617 |
| 4XCU | Pyridyl acetic acid | -4.71 | 1 | ARG559 |
| 4XCU | Eicosatetraynoic acid | -6.12 | 4 | HIS638, ASP641, LYS644 |
| 4XCU | Promethazine<br>sulfoxide | -5.24 | - |  |
| 4XCU | Sulfadoxine | -5.89 | 4 | ASN557, SER647, ASP612,<br>ARG559 |
| 4ASD | Trigonelline | -3.89 | 2 | ARG932 |
| 4ASD | Salicylamide | -4.41 | 3 | SER1104, GLU1097, ASP1058 |
| 4ASD | Pyridyl acetic acid | -3.88 | 2 | ARG932 |
| 4ASD | Eicosatetraynoic acid | -1.83 | 1 | ARG932 |
| 4ASD | Cyclopenin | -4.86 | 3 | ARG932, SER1104. ALA1127 |
| 4ASD | Promethazine<br>sulfoxide | -4.8 | - |  |
| 4ASD | Sulfadoxine | -4.9 | 3 | ARG932, SER1104. ALA1127 |
| 1SVC | Trigonelline | -4.18 | 2 | LEU45, GLU233 |
| 1SVC | Salicylamide | -3.62 | 2 |  |
| 1SVC | Pyridyl acetic acid | -4.76 | 2 | ARG255 |
| 1SVC | Eicosatetraynoic acid | -4.87 | 2 | HIS307, ARG308 |
| 1SVC | Cyclopenin | -5.32 | 3 | ILE283, LYS337 |
| 1SVC | Promethazine<br>sulfoxide | -5.36 | - | - |
| 1SVC | Sulfadoxine | -4.3 | 3 | ASP305, HIS307, THR304 |

## Acknowledgement

**Nil**

